# ATP induces protein folding, inhibits aggregation and antagonizes destabilization by effectively mediating water-protein-ion interactions, the heart of protein folding and aggregation

**DOI:** 10.1101/2020.06.21.163758

**Authors:** Jian Kang, Liangzhong Lim, Jianxing Song

## Abstract

Many, particularly β-dominant proteins, are prone to misfolding/aggregation in the crowded cells, a hallmark of ageing and neurodegenerative diseases including ALS. ATP provides energy to drive supramolecular machineries to control protein hemostasis in modern cells. Recently ATP was decoded to hydrotropically inhibit/dissolve liquid-liquid phase separation (LLPS) and aggregation/fibrillation at millimolar concentrations. We also found that by specific binding, ATP induces and subsequently dissolves LLPS, as well as inhibits fibrillation. Nevertheless, no report shows that ATP can directly induce protein folding. Here, by selecting two aggregation-prone ALS-causing proteins with the unfolded states, we successfully visualized the effects of ATP and 11 molecules with NMR directly on their folding and aggregation. The study reveals for the first time that ATP can induce folding at molar ratios of 2-8, the highest efficiency known so far. Intriguingly, this inducing-capacity comes from triphosphate, a key intermediate in prebiotic chemistry, which, however, also triggers aggregation. Most unexpectedly, upon joining with adenosine, the ability of triphosphate to trigger aggregation is shielded. Marvelously, ATP emerged to manifest three integrated abilities: to induce folding, inhibit aggregation and increase stability, that are absent in ATPP, AMP-PCP and AMP-PNP. Our study sheds the first light on previously-unknown roles of ATP in energy-independently controlling protein folding and aggregation by effectively mediating water-protein-ion interactions. Therefore, ATP might be not just irreplaceable for solving protein folding and aggregation problems simultaneously in primitive cells for Origin of Life, but also energy-independently operating in modern cells to regulate protein homeostasis fundamentally critical for physiology and pathology.

---

Many proteins need to fold from the unfolded state (U) into the marginally stable native state (N), but the high-resolution mechanism still remains a mystery (1-10). Cell is extremely crowded with proteins (>100 mg/ml), and consequently many, particularly β-sheet-dominant proteins, are prone to misfolding/aggregation, a pathological hallmark of ageing and neurodegenerative diseases including amyotrophic lateral sclerosis (ALS) (8-22). Modern cells handle protein folding and aggregation problems by supramolecular machineries energetically driven by ATP (9), the universal energy currency but mysteriously with very high concentrations (2-12 mM). Recently, ATP was decoded to hydrotropically inhibit/dissolve liquid-liquid phase separation (LLPS) and aggregation/fibrillation at the proteome-wide scale (12-14). We further discovered that by specific binding, ATP induces and subsequently dissolves LLPS, as well as inhibits aggregation/fibrillation (16-18). Nevertheless, so far, no report shows that ATP can directly induce protein folding.

Here by high-resolution multidimensional NMR, we successfully visualized the effects of ATP and 11 molecules (Fig. S1) directly on the folding reactions and aggregation of two unique ALS-causing proteins. The study decoded for the first time that ATP has the capacity in inducing protein folding with the highest efficiency known so far, which unexpectedly comes from triphosphate, a key intermediate in prebiotic evolution (23). However, triphosphate owns the strong ability to trigger aggregation. Marvelously, by joining adenosine only to triphosphate, nor tetraphosphate, nor methylenediphosphonate, nor imidodiphosphate, ATP emerged to acquire three integrated abilities apparently by optimizing the interplay of the water-attracting triphosphate chain and hydrophobically-interacting purine ring: to induce folding, inhibit aggregation and increase stability via mediating water-protein-ion interactions. Therefore, ATP might be not just irreplaceable for Origin of Life by simultaneously solving protein folding and aggregation problems in primitive cells, but still energy-independently operating in modern cells to control protein homeostasis to antagonize diseases and ageing.

## Selection of the model proteins

In 1960s, Anfinsen decrypted the principle for protein folding on RNase A, which universally holds for all proteins regardless of their secondary and tertiary structures, including a list of well-studied model proteins (Fig. S2). However, the relationship between protein folding, aggregation and thermodynamic stability is extremely complex, and it still remains a fundamental challenge to dissect them (1-10).

Therefore, here we aimed to overcome this complexity by visualizing the effects of ATP and 11 molecules (Fig. S1) directly on the folding reactions and aggregation relevant to human diseases. In this regard, the model proteins should satisfy the following criteria: 1) well-studied human proteins of >100 residues with complex and diverse tertiary topologies composed of all secondary structure elements, whose aggregation is relevant to human diseases; 2) most importantly, existence of the unfolded state without any denaturants; 3) no specific/significant interaction with nucleic acids or/and nucleotides to exclude the possibility that the effects of ATP and its analogues may result from the specific binding.

Exhaustive studies have revealed that unlike β-sheet-dominant proteins whose structures are mainly stabilized by long-range interactions, helix-dominant proteins stabilized by local interactions including T4 lysozyme (6) and myoglobulin (7) would not be rendered into the unfolded state by slight sequence variations under non-denaturing conditions. For example, even the deletion of all four disulfide bridges only rendered α-lactalbumin into the molten-globule state which still retains both native-like secondary structures and tertiary topology but becomes unamendable for NMR studies (24).

Strikingly, ALS-causing C71G mutant of human profilin-1 (C71G-hPFN1) and nascent superoxide dismutase 1 (hSOD1) satisfy all three criteria. hPFN1 is a well-studied model protein of 140 residues with pI of 8.6, which adopts a seven stranded antiparallel β-sheet sandwiched by α-helices composed of 45% α−helix, 32% β−strand and 23% turn/loop. No specific/significant interaction with nucleic acids or nucleotides is required for its functions. Our previous NMR characterization also showed that unlike TDP-43 (17) and FUS (18) RRM domains, the WT-hPFN1 has no specific/significant interaction with ATP (18). Importantly, C71G-hPFN1 (Fig. 1A) characteristic of severe aggregation both *in vitro* and *in vivo* is the most toxic mutant causing ALS by gain of toxicity (21). Our extensive CD and NMR studies on the conformations, dynamics and aggregation showed that C71G-hPFN1 is in equilibrium between the folded and unfolded states without any denaturants, and prone to aggregation in high-salt buffers, thus mimicking ALS-causing aggregation in the cellular environments with ∼150 mM salts (19).

**Fig. 1.**
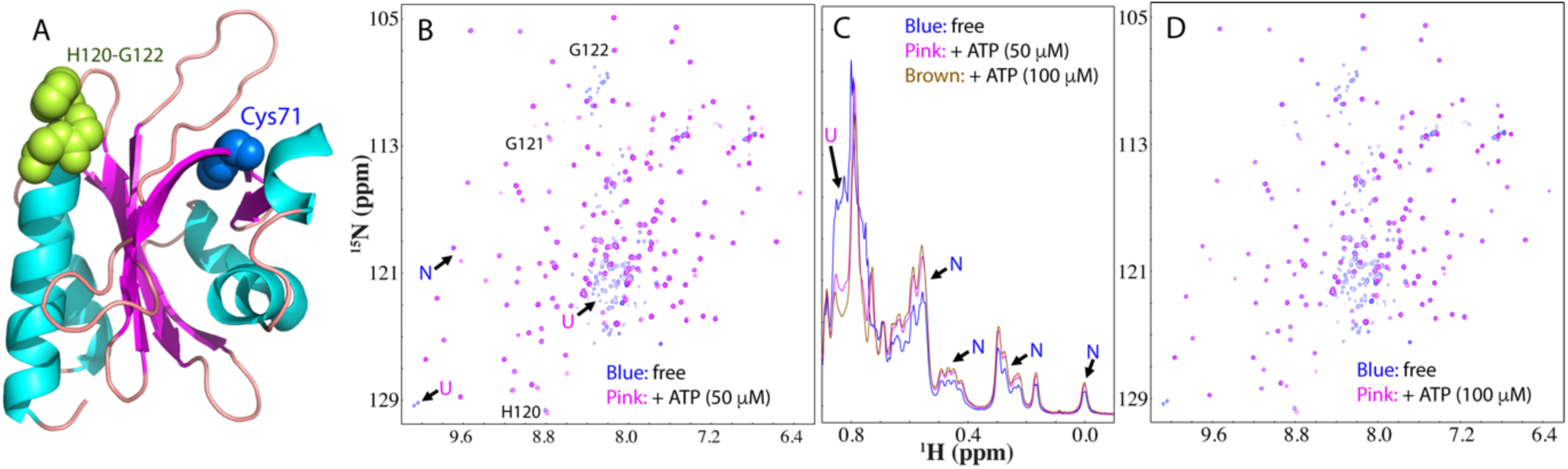
ATP shifts the conformational equilibrium to favor the folded state. (A) The three-dimensional structure of hPFN1 (PDB ID of 2PAV) with Cys71 displayed in blue spheres, and His120-Gly121-Gly122 in yellow-green spheres. (B) Superimposition of HSQC spectra of ^15^N-labeled C71G-PFN1 at a concentration of 50 μM in the absence (blue) and presence of ATP at 50 μM (pink). (C) Up-field 1D NMR spectra of C71G-PFN1 in the presence of ATP at different concentrations. (D) Superimposition of HSQC spectra of C71G-PFN1 in the absence (blue) and in the presence of ATP (pink) at a molar ratio of 1:2. Some characteristic NMR signals of the folded native (N) and unfolded (U) states were indicated by arrows. The assignments are labeled for three HSQC peaks with ATP-induced shifts,

The 153-residue hSOD1 with pI of 4.5 has been exhaustively studied before for both physiological functions and pathological roles (10,11,19,22). The native hSOD1 with the zinc, copper and disulfide bridge Cys57-Cys146 adopts a β-barrel composed of eight antiparallel β-strands arranged in a Greek key motif consisting of 20% α−helix, 51% β−strand and 29% turn/loop. Moreover, it also has no interaction with nucleic acids or/and nucleotides. Since 1993, >180 mutations have been identified to lead to ALS. In particular, the nascent hSOD1 without any mutation but lacking metal ions and disulfide bridge is also prone to aggregation *in vitro* and *in vivo*, thus initiating ALS (20,22). Previously we conducted extensive NMR and CD studies on the conformations, dynamics and aggregation and showed that the nascent hSOD1 is completely unfolded without any stable secondary and tertiary structures (20).

## ATP induces folding of the unfolded state of C71G-hPFN1

We first studied the effect of ATP on C71G-hPFN1 co-existing between the folded and unfolded states (19), as evidenced by the presence of two set of NMR HSQC peaks (Fig. S3A). Here we successfully achieved NMR assignments of both WT-hPFN1 and C71G-hPFN1 except for some missing or overlapping peaks. WT-hPFN1 and the folded state of C71G-hPFN1 have highly similar (ΔCα-ΔCβ) values (Fig. S4A), indicating that they have very similar conformations. By contrast, the absolute values of (ΔCα-ΔCβ) of the unfolded state are much smaller than those of its folded state (Fig. S4B), clearly indicating that the unfolded state is highly disordered without any stable secondary and tertiary structures.

Strikingly, when ATP was added even only at a molar ratio of 1:0.5 (C71G:ATP), the intensity of HSQC peaks of the unfolded state became reduced while those of the folded state slightly increased (Fig. S3B and S5A). The 1D peak intensity of the methyl group from the unfolded state also became reduced while those of the folded state slightly increased (Fig. S3C). When ATP was added to 1:1, the HSQC peak intensity of the unfolded state became further reduced while those of the folded state increased (Fig. 1B, 1C and S5B). At 1:2, HSQC peaks of the unfolded state became completely disappeared (Fig. 1D), and the 1D peak intensity of the methyl group of the folded state increased (Fig. 1C). Further addition of ATP to 1:20 (1 mM) showed no significant change of HSQC spectrum (Fig S3D). These results indicate that ATP is capable of completely converting the unfolded state into the folded state at a very low molar ratio. It is also worth to note that ATP induced no significant shift of HSQC peaks of the unfolded state, and only some minor shifts of HSQC peaks of His120, Gly121 and Gly122 of the folded state (Fig. 1A and 1B), exactly as we previously observed on WT-hPFN1 (17). This suggests that ATP has no specific/significant interaction with both states of C71G-hPFN1. We also increased the C71G-hPFN1 concentration to 100 μM, and the ATP concentration required to completely convert the unfolded state doubled (200 μM). This suggests that the induction by ATP is dependent of the molar ratio between C71G-hPFN1 and ATP.

## The inducing capacity of ATP comes from its triphosphate chain

To determine the group of the ATP molecule responsible for this inducing capacity, we subsequently titrated C71G-hPFN1 with a series of related molecules (Fig. S1). For ADP, only at 1:8 (C71G:ADP), HSQC peaks of the unfolded state became completely disappeared and interestingly, most HSQC peaks with ADP at 1:8 are almost completely superimposable to those with ATP at 1:2 (Fig. S6). This indicates that ADP still has the capacity in shifting the equilibrium but becomes much weaker than ATP. We also titrated with AMP (Fig. S7), but even with the concentrations up to 20 mM (1:400), AMP was still unable to completely convert the unfolded state into the folded state. We further titrated with adenosine and even at the highest concentrations of 5 mM due to its low solubility, no significant change was detected for HSQC spectra (Fig. S8). This reveals that adenosine also has no detectable binding to the protein.

The results strongly suggest that the capacity of ATP in inducing protein folding comes from its triphosphate chain. Indeed, when titrated with sodium triphosphate (PPP), at 1:1 (C71G:PPP) the HSQC peak intensity of the unfolded state also became significantly reduced (Fig. 2A and S9). Furthermore, the 1D peak intensity of the methyl group of the unfolded state also became reduced while those of the folded state slightly increased (Fig 2B). At 1:2, HSQC peaks of the unfolded state became completely disappeared (Fig. 2C), and the intensity of the methyl group of the folded state further increased (Fig. 2B). Very strikingly, C71G-hPFN1 with PPP at 1:2 has both HSQC (Fig. 2D) and 1D (Fig. 2B) spectra very similar to those with ATP at 1:2. On the other hand, completely different from ATP which induces no aggregation even at 20 mM, further addition of PPP (>0.15 mM) led to the visible precipitation and NMR signals become too weak to be detectable. The results reveal that the capacity of ATP in inducing folding indeed comes from its triphosphate chain but the isolated PPP owns a very strong ability to induce aggregation of C71G-hPFN1 with significant packing defects most likely due to the electrostatic screening effect (15).

**Fig. 2.**
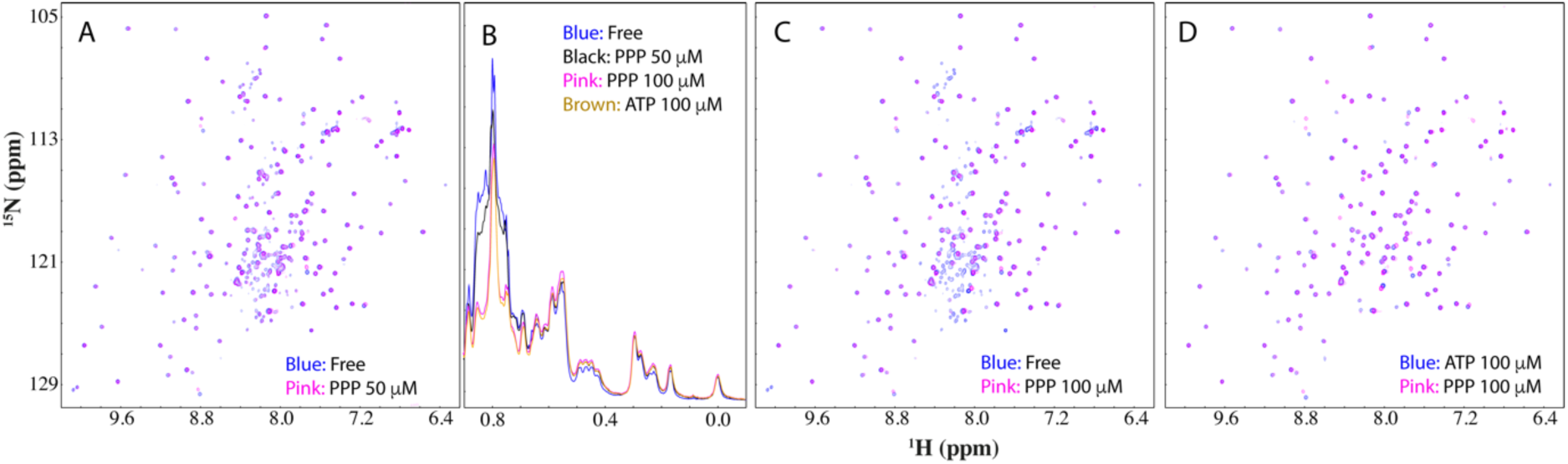
Triphosphate (PPP) shifts the conformational equilibrium to favor the folded state. (A) Superimposition of HSQC spectra of ^15^N-labeled C71G-PFN1 at a concentration of 50 μM in the absence (blue) and in the presence of PPP (pink) at a molar ratio of 1:1. (B) Up-field 1D NMR spectra of C71G-PFN1 in the presence of PPP at different ratios. (C) Superimposition of HSQC spectra of C71G-PFN1 in the absence (blue) and in the presence of PPP (pink) at a molar ratio of 1:2 (D) Superimposition of HSQC spectra of C71G-PFN1 in the presence of ATP at 1:2 (blue) and PPP at 1:2 (pink).

We further titrated with sodium salts of diphosphate (PP), phosphate (P) and chloride. Interestingly, PP has an inducing capacity very similar to ADP: only at 1:8 (C71G:PP), HSQC peaks of the unfolded state became completely disappeared (Fig. S10). Again, completely different from ADP which induced no precipitation even at 20 mM (C71G:ADP=1:400), PP started to induce visible precipitation at concentrations higher than 0.4 mM. Moreover, sodium phosphate and chloride failed to convert the unfolded state into the folded state with the concentration up to 5 mM for sodium phosphate (Fig. S11), and 10 mM for NaCl (Fig. S12), where NMR samples started to show visible precipitation. Noticeably, the addition of three salts appeared to induce the intensity reduction of both folded and unfolded peaks most likely due to salt-induced aggregation as we previously characterized (15,19,20). These results imply that the inducing capacity of PPP might not be simply resulting from its high ionic strength because the ionic strength of sodium salts of triphosphate and diphosphate are only 15-time and 10-time respectively stronger than sodium chloride while only 2.5-time and ∼1.67-time respectively stronger than sodium phosphate (12).

We further assessed the effects of three ATP analogues, namely Adenosine 5’-(pentahydrogen tetraphosphate) (ATPP), Adenylyl-imidodiphosphate (AMP-PNP) and Methyleneadenosine 5-triphosphate (AMP-PCP) (Fig. S1). Very unexpectedly, ATPP induced the complete conversion of the unfolded state into the folded state only at 1:2 (C71G:ATPP), which is very similar to ATP (Fig. S13A). This result indicates that the inclusion of one extra phosphate failed to significantly enhance the capacity of ATP to induce folding. Moreover, ATPP showed significant ability to trigger the aggregation of C71G-hPFN1 as evidenced by the fact that at 1:3 (C71G:ATPP), the C71G-hPFN1 protein started to aggregate and many HSQC peaks became too weak to be detected. Further addition of ATPP immediately triggered visible precipitation and no NMR signal could be detected.

Furthermore, upon replacing the oxygen atom linking the β− and γ−phosphorous atom with carbon atom, the capacity of AMP-PCP to induce folding significantly reduced to the level very similar to that of ADP: only at a ratio of 1:8 (C71G:AMP-PCP) the unfolded state completely transformed into the folded state (Fig. S13B). On the other hand, unlike ADP which triggered no precipitation even at 20 mM, upon adding AMP-PCP with concentrations > 1 mM, C71G-hPFN1 also started to precipitate after 10 minutes. Intriguingly, AMP-PNP with this oxygen atom replaced by nitrogen atom has the capacity in inducing folding very similar to that of ADP and AMP-PNP (Fig. S13C). However, its ability to trigger aggregation is even higher than that of AMP-PCP: at 1:10 (C71G:AMP-PNP), C71G-PFN1 started to precipitate after 15 minutes.

## ATP induces the folding of the unfolded nascent hSOD1

To assess whether the folding-inducing capacity of ATP and PPP is only specific for C71G-hPFN1, or general for other proteins with different functions, physicochemical properties and structures, we subsequently studied their effects on the nascent hSOD1 (Fig. 3A). As we previously characterized (19), the nascent hSOD1 is highly unfolded, which has HSQC (Fig. S14A) and 1D (Fig. S14B) spectra typical of an unfolded protein (7). Upon adding ATP to 1:2, no significant change was observed on HSQC (Fig. S14C) and 1D spectra (Fig. S14B). Nevertheless, when the ratio was increased to 1:4, well-dispersed HSQC peaks and very up-field 1D signals manifested, which was saturated at 1:8 (Fig. 3B, 3C and S15), indicating that the folded state was formed. Further addition of ATP to 1:4000 (20 mM) only induced some minor shifts of HSQC peaks (Fig S14D), indicating that unlike C71G-hPFN1, ATP could not completely convert the unfolded state of hSOD1 into the folded state.

**Fig. 3.**
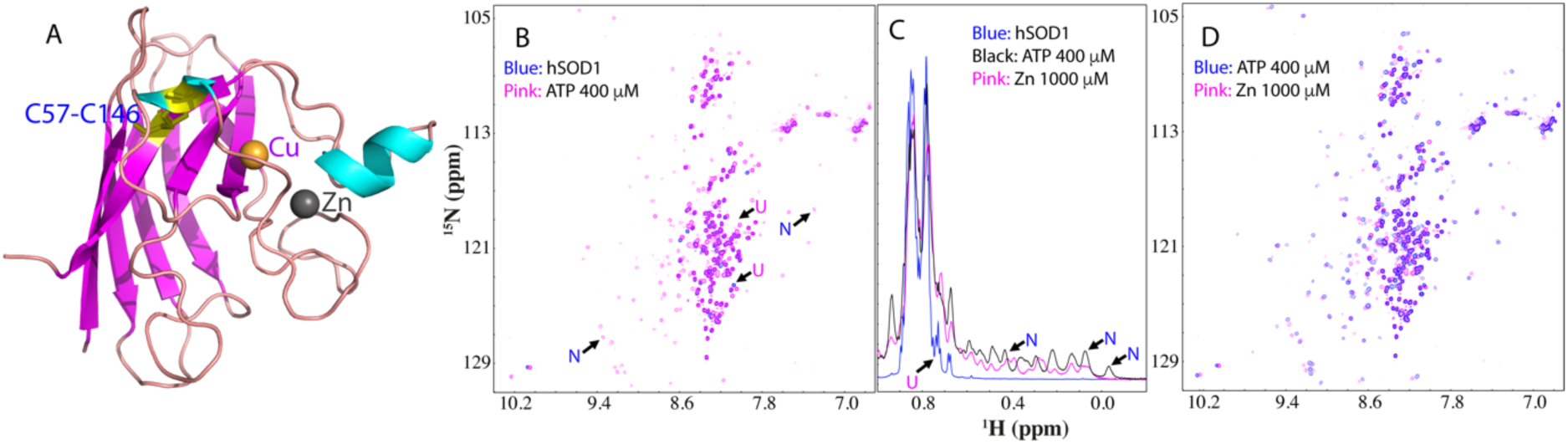
ATP induces the folding of the unfolded nascent hSOD1. (A) The three-dimensional structure of hSOD1 (PDB ID of 1MFM) with the cofactors zinc and copper cations displayed in spheres and disulfide bridge Cys57-Cys146 in yellow sticks. (B) Superimposition of HSQC spectra of ^15^N-labeled hSOD1 at a concentration of 50 μM in the absence (blue) and in the presence of ATP (pink) at a molar ratio of 1:8. (C) Up-field 1D NMR spectra of hSOD1 at 50 µM in the absence (blue) and in the presence of ATP at 1:8 (pink) and zinc cation at 1:20 (black). (D) Superimposition of HSQC spectra of hSOD1 in the presence of ATP at 1:8 (blue) and in the presence of zinc cation at 1:20 (pink).

Interestingly, many HSQC peaks of the folded states induced by ATP here and by zinc we previously reported (20) are superimposable (Fig. 3D), while 1D signals have some differences (Fig. 3C), implying that they have similar backbone conformations but some difference in sidechain packing (19). We also found that like ATP, PPP could induce the folding of hSOD1. At 1:2, no significant change was observed on HSQC (Fig. S16A) and 1D (Fig. S16B) spectra. However, at 1:4, the folded population was formed as evidenced by the manifestation of the well-dispersed HSQC and up-field 1D peaks and this was also saturated at 1:8 (Fig. S16B, S16C and S17). Very interestingly, both HSQC (Fig. S16D) and 1D (Fig. S16B) spectra with PPP at 1:8 are very similar to those with ATP at 1:8, indicating the high similarity of the conformations induced by ATP and by PPP. As also observed on C71G-hPFN1, further addition of PPP also triggered severe aggregation of the nascent hSOD1.

Previously we showed that the unfolded nascent hSOD1 could only be significantly induced to fold by its cofactor zinc and it became saturated at 1:20. However, even at 1:20, the unfolded state still co-existed with the folded state, and we have successfully obtained their residue-specific NMR conformations and dynamics (20). Furthermore, we screened 11 other cations including Na^+^, K^+^, Ca^2+^, Mg^2+^, Cu^2+^, Fe^2+^ and Al^3+^ as well as Co^2+^, Ni^2+^, Cd^2+^, and Mn^2+^ which are commonly used to replace zinc in the native SOD1 for various structural studies, but all of them failed to significantly induce the folding of hSOD1 even at 1:40 (Fig. S18) where the samples started to precipitate. Very unexpectedly, ATP and PPP not only can induce folding of hSOD1 as zinc, but also have much higher capacity than zinc.

Therefore, we set to examine whether ATP and zinc together can completely shift the unfolded state. We added ATP to hSOD1 with the pre-existence of zinc at 1:20 (Fig. S19A and S19B). However, even with extra addition of ATP at 1:8, no significant change was observed for both HSQC (Fig. S19C and S19D) and 1D spectra (Fig. S19B), indicating that ATP and Zn^2+^ together are still unable to completely convert the unfolded state into the folded state.

## TMAO fails to induce folding at concentrations that trigger aggregation

Previously, the best-known small molecule with the general capacity in inducing protein folding is the natural osmolyte, trimethylamine N-oxide (TMAO) (Fig. S1) (3,25), which was shown to induce the folding of the reduced RNase T1 with the co-existence of the unfolded and folded states at 10 μg/ml with TMAO concentrations at ∼1.2 M (25). Here we first titrated C71G-hPFN1 with TMAO and the results showed that even at 1:20 with a TMAO of 1 mM, no significant change were observed for both the up-field 1D (Fig. S20A) and HSQC (Fig. S20B) spectra. At 1:200, for both folded and unfolded states, the intensity of 1D peaks reduced (Fig. S20A), while HSQC peaks showed some shifts (Fig. S20C). However, no significant disappearance of HSQC peaks from the unfolded state was observed even with TMAO concentration up to 50 mM (Fig. S20D). Further addition of TMAO triggered visible precipitation and consequently no NMR signal could be detected.

Subsequently we titrated the nascent hSOD1 with TMAO and the results showed that even at 1:200 with a TMAO concentration of 10 mM, no significant change were observed for both the up-field 1D (Fig. S21A) and HSQC (Fig. S21B) spectra. At 1:1000, for both folded and unfolded states, the intensity of 1D spectra reduced (Fig. S21A), while HSQC peaks have significant shifts (Fig. S21C). Nevertheless, no detectable formation of the folded state was observed as judged by 1D and HSQC spectra, even with TMAO concentration up to 100 mM (Fig. S21D). Further addition of TMAO also triggered visible precipitation of the protein.

The results with TMAO here suggest that consistent with the previous studies (25), TMAO needs much higher concentrations than ATP to induce folding. However, for aggregation-prone C71G-PFN1 and nascent hSOD1, the polar TMAO is also capable of triggering aggregation even at concentrations much lower than those required for inducing protein folding.

## ATP induces protein folding by enhancing the intrinsic folding propensity

So why could ATP completely convert the unfolded state of C71G-hPFN1 but failed for hSOD1? Previously, we have collected HSQC-NOESY spectrum of the hSOD1 sample with zinc at 1:20, and found that no NOE cross peaks manifested for the exchange between the folded and unfolded states, indicating that for hSOD1 with zinc cation, the exchange of the two states is much slower than the NMR time scale. In other words, the energy barrier separating the folded and unfolded states of hSOD1 are large (20). Indeed, it was shown that the complete formation of the native hSOD1 needs further copper-load and covalent reaction to form disulfide bridge that can only be catalyzed by human copper chaperone for SOD1 (hCCS) (22).

Here we collected HSQC-NOESY spectrum for C71G-hPFN1, and subsequently found the presence of the cross peaks for the exchange between the folded and unfolded states (Fig. S22). Therefore, by using the well-established NMR methods (26,27), we successfully mapped out the populations of 55.2% and 44.8% respectively for the folded and unfolded states, and the exchange rate of ∼11.7 Hz (Table S1). Moreover, we also determined the NMR dynamics on the ps-ns time scale by acquiring ^15^N backbone relaxation data T1, T2 and hNOE for both WT-hPFN1 and C71G-hPFN1 (Fig. S23), and subsequently performed the “model-free” analysis, which generates squared generalized order parameters, S^2^, reflecting the conformational rigidity on ps-ns time scale. S^2^ values range from 0 for high internal motion to 1 for completely restricted motion in a molecular reference frame (28,29). As shown in Fig 4A, the majority of the WT-hPFN1 residues has S^2^ > 0.76 (with the average value of 0.89), suggesting that WT-hPFN1 has very high conformational rigidity (Fig. 4B). By contrast, many residues of the folded state of C71G-hPFN1 have S^2^ < 0.76 (Fig. 4A) (with the average value of 0.73) (Fig. 4C), revealing that even the folded state of C71G-hPFN1 becomes more dynamic than WT-hPFN1 on ps-ns time scale. Furthermore, the overall rotational correlation time (τc) of WT-hPFN1 was determined to be 7.5 ns while that of C71G-hPFN1 was 7.8 ns, implying that C71G-hPFN1 becomes less compact, completely consistent with their translational diffusion coefficients (∼1.12 ± 0.03 × 10^−10^ m^2^/s for WT-hPFN1, and ∼1.03 ± 0.02 × 10^−10^ m^2^/s for C71G-hPFN1).

**Fig. 4.**
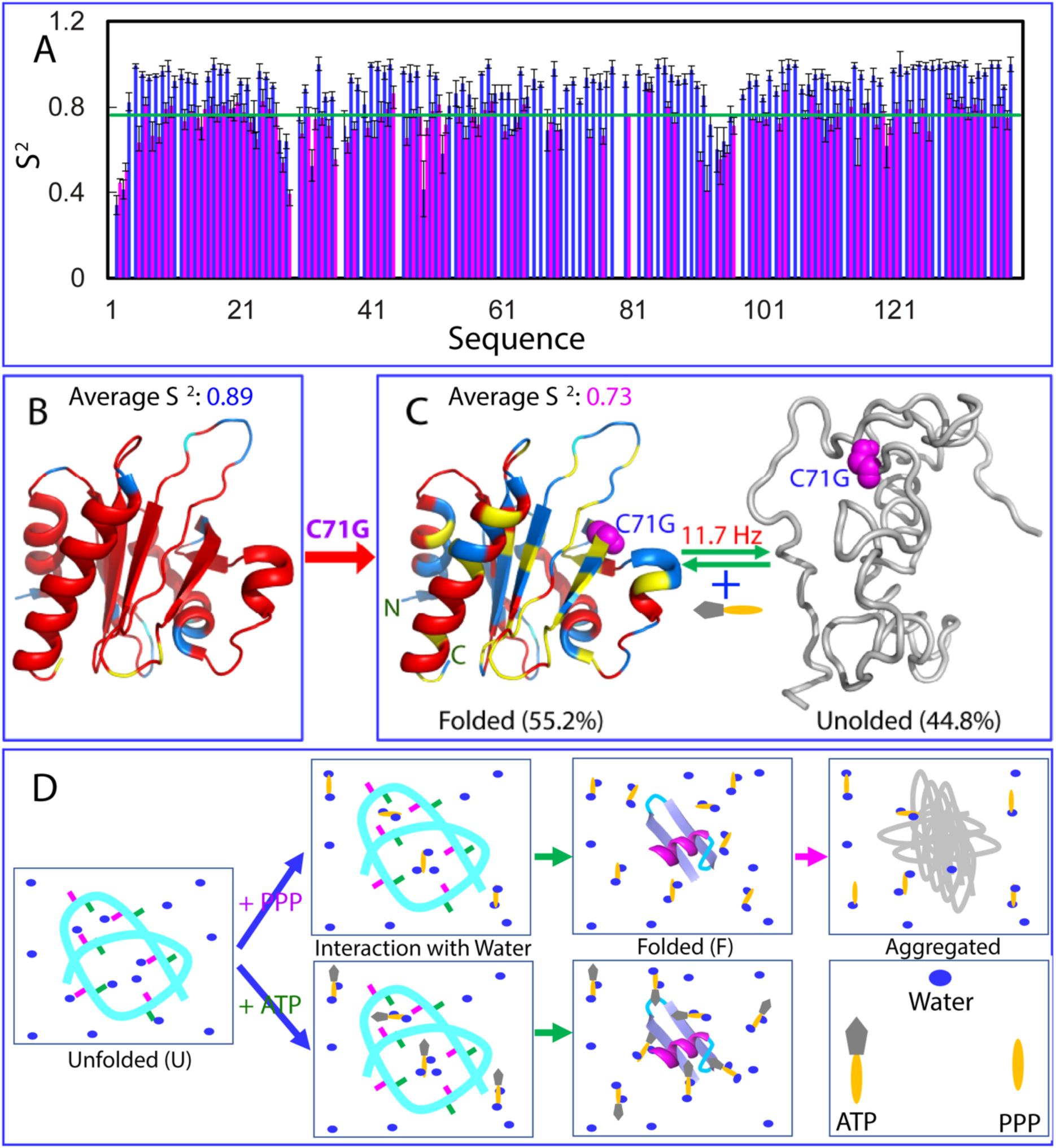
A speculative model for ATP/PPP to modulate protein folding and aggregation. (A) Generalized squared order parameter (S^2^) of WT-hPFN1 (blue) and the folded state of C71G-hPFN1 (pink). The green line has a S^2^ value of 0.76 (average – STD). (B) The structure of hPFN1 (PDB ID of 2PAV) with S^2^ values (with the average value of 0.89) of WT-hPFN1 mapped on. (C) A diagram to show the co-existence of the folded (with the average S^2^ value of 0.73) and unfolded states of C71G-hPFN1. S^2^ values are also mapped onto the folded state of C71G. Cyan is used for indicating Pro residues, and the yellow for residues with missing or overlapping HSQC peaks. Red is for residues with S^2^ values > 0.76 and blue for residues with S^2^ values < 0.76. The folded state with a population of 55.2% and unfolded state with a population of 44.8% have been experimentally mapped out to undergo a conformational exchange at 11.7 Hz. (D) A speculative model for ATP/PPP to mediate protein folding and aggregation. For the unfolded state (U), many backbone atoms are hydrogen-bonded with water molecules. ATP/PPP can use the triphosphate chain to effectively attract water molecules out from hydrogen-bonding with the backbone atoms, thus favoring the formation of the folded state (F). However, in the presence of the isolated triphosphate with highly negative charges that produce significant electrostatic screening effects, a protein exemplified by C71G-PFN1 with significant exposure of hydrophobic patches resulting from the defects in tertiary packing will become severely aggregated. By contrast, ATP can not only utilize its triphosphate chain to induce folding as effective as PPP, but simultaneously use its aromatic ring to dynamically interact with the exposed hydrophobic patches, consequently preventing aggregation, as well as increasing the thermodynamic stability of the protein.

The results imply that ATP induces protein folding by enhancing the intrinsic folding capacity already encoded by the protein sequence, and consequently it failed to induce the complete conversion of the unfolded population of hSOD1 because this conversion needs the covalent formation of the disulfide bridge (20,22).

## ATP enhances thermodynamic stability

We also measured the thermal stability by differential scanning fluorimetric (DSF) method we previously utilized to assess the effect of ATP on other proteins (17,18). Interestingly, WT-hPFN1 at 10 μM has a melting temperature (Tm) of ∼56 degree while the addition of ATP even up to 20 mM triggered no significant change (Fig. S24A). By contrast, C71G-hPFN1 at 10 μM without ATP has no cooperative unfolding signal (Fig S24B), likely due to the absence of tight tertiary packing or/and co-existence of two states (19,30). Interestingly in the presence of ATP of 20 μM (a molar ratio of 1:2), a cooperative unfolding signal was observed with Tm of 32 degree. Addition of ATP to 1 mM (1:100) led to the increase of Tm to 38 degree, and further increase to 40 degree at 20 mM (Fig. S24C). By contrast, we performed DSF measurements of C71G-hPFN1 with addition of PPP and ATPP at various ratios and only obtained unfolding curves with no cooperative unfolding signals but high noises, implying that although ATP, PPP and ATPP have similar capacity in inducing folding, PPP and ATPP failed to enhance tight tertiary packing of the folded state or/and triggered dynamic aggregation even before the visible precipitation. Indeed, previously we found that a protein could still have native-like NMR spectra although its tight packing was disrupted to different degrees (30).

## Mechanisms for ATP to induce folding, inhibit aggregation and enhance stability

The present study decoded for the first time that ATP, the universal “energy currency”, also has the capacity to energy-independently induce protein folding at very low ratios of 2-8, which can be satisfied in most, if not all cells. Unexpectedly, this capacity of ATP comes from triphosphate, a key intermediate for various prebiotic chemical reactions to generate building units for constructing primitive cells, thus allowing Origin of Life (23). However, triphosphate strongly triggers aggregation of proteins with the deficient packing. Marvelously the formation of ATP by linking to adenosine not only retains the capacity of triphosphate to induce folding, but surprisingly shields its ability to trigger aggregation. Moreover, ATP becomes even capable of inhibiting aggregation/fibrillation at high concentrations (12-17). As compared to TMAO that induced no detectable folding of C71G-hPFN1 and nascent hSOD1 respectively at 50 and 100 mM, which are sufficient for triggering severe aggregation, ATP is thus established to induce folding with the highest efficiency known so far.

In the context of inducing protein folding, triphosphate appears to be optimized for both chain length and structure, because: 1) while the reduction of the chain length in ADP and AMP significantly decreases the inducing capacity, the increase in ATPP also fails to enhance the capacity; 2) slight variations of the chemical structures in AMP-PNP and AMP-PCP, which have been commonly utilized to mimic ATP in various studies, significantly reduces the inducing capacity. With regard to aggregation, triphosphate, diphosphate and phosphate all have the ability to trigger aggregation. Surprisingly, however, upon joining with adenosine to form ATP, ADP and AMP, the ability to trigger aggregation becomes shielded. Intriguingly, however, the variations of the chemical structures of the triphosphate chains in ATPP, AMP-PNP and AMP-PCP lead to dramatic disruption of the shielding, and consequently unlike ATP, these analogues are still able to trigger aggregation.

So, what could be the mechanism for ATP/triphosphate to induce folding at such low molar ratios for the proteins of unrelated functions and opposite pI values as well as different secondary structure contents and tertiary topologies but without specific/significant interaction? In principle, a molecule can mediate folding of a protein by interacting with its unfolded or/and folded states, or/and solvation/hydrogen-bonding with water molecules (1-10,15,25,31). Previously, the hydrogen-bonding of water molecules with proteins, particularly with the backbone atoms, was shown to play a key role in protein folding (2-4,25). The unfolded state (U) will be favored if the backbone atoms are highly hydrogen-bonded with water molecules, while the native state (N) will be favored if the backbone atoms are shifted to form intramolecular hydrogen bonds (3). Indeed, the polar TMAO induces folding by interacting with water molecules to disrupt the backbone-water hydrogen bonds so as to facilitate the formation of intramolecular hydrogen bonds (3,25). As the backbone structures and solvation are common to all proteins, TMAO thus owns the general ability to induce folding of proteins regardless of their secondary and tertiary structures (3,25).

As ATP induces protein folding without significant/specific interaction with both unfolded and folded states, here we propose that ATP utilizes its triphosphate chain to induce folding by the same mechanism for TMAO but with the much higher efficiency, as ATP/triphosphate has the very high capacity in interacting with water molecules over protein surfaces (12-14,16,31). As illustrated by a speculative model (Fig. 4D), ATP/triphosphate can efficiently attract water molecules out from hydrogen-bonding or/and hydration shell with the protein backbone atoms. Consequently, the backbone atoms will be shifted into forming the intramolecular hydrogen bonds to favor folding from the unfolded (U) into the folded (F) states. Therefore, like TMAO (3,25), ATP/PPP can also induce folding of proteins with diverse physicochemical properties, secondary and tertiary structures.

On the other hand, however, a protein exemplified by C71G-PFN1 with a significant exposure of hydrophobic patches will be also induced to aggregate by triphosphate, which can significantly impose the screening effect (10,15). Amazingly, upon joining with adenosine, ATP can use its hydrophobic aromatic ring to be dynamically clustered over the hydrophobic patches of a protein and thus protecting them from directly contacting bulk solvent, which consequently acts to shield the aggregation-inducing ability of triphosphate. However, due to the extremely weak interactions, only at millimolar concentrations, ATP manifests the ability to increase the thermodynamic stability by enhancing the tertiary packing (12-14,17). Therefore, unlike TMAO which only has polar groups to induce folding by interacting with water molecules, ATP appears to represent the optimized combination of the hydrophobically-interacting purine ring and water-attracting triphosphate chain to achieve the extremely effective mediation of water-protein-ion interactions, which manifests the three integrated abilities. This finding thus reveals a novel principle to further engineer three key abilities into one molecule by optimizing the interplay of the hydrophobically-interacting and water-attracting groups, which should hold promising potential for developing therapeutic molecules as well as other applications.

Our current study also implies that nature selects triphosphate as the key molecule for prebiotic evolution (23), also because of its extremely high efficiency in inducing protein folding. Nevertheless, only the emergence of ATP solved protein folding and aggregation problems simultaneously. With our new results added, ATP has now been established to modulate both sides of protein homeostasis: induce folding and inhibit/dissolve aggregation also by various energy-independent mechanisms (12-17,31). Therefore, ATP might play a central role in simultaneously solving protein folding and aggregation problems in primitive cells which lacked any modern ATP-energy-driven machineries, and consequently be irreplaceable for Origin of Life. It is also anticipated that even in modern cells, ATP is still energy-independently operating at fundamental levels to control protein homeostasis together with the well-studied ATP-energy-dependent chaperone and disaggregase machineries. The integrated capacity of ATP might be particularly important for handling the aggregation-prone proteins exemplified by C71G-hPFN1 with the intrinsically defective packing due to genetic mutations, which thus needs a ubiquitous availability of ATP. This may explain the long-standing fact that the risk of neurodegenerative diseases dramatically increases with being aged because in aged cells ATP concentrations are significantly reduced.

## Acknowledgment

This study is supported by Ministry of Education of Singapore MOE Tier 1 grants R-154-000-B45-114 and R-154-000-B92-114 to Jianxing Song..

